# Evidence of Müller glia conversion into retina ganglion cells using Neurogenin2

**DOI:** 10.1101/399717

**Authors:** Roberta Pereira de Melo Guimarães, Bruna Soares Landeira, Diego Marques Coelho, Daiane Cristina Ferreira Golbert, Mariana S. Silveira, Rafael Linden, Ricardo A. de Melo Reis, Marcos R. Costa

## Abstract

Macular Degeneration, Glaucoma, and Retinitis Pigmentosa are all leading causes of irreversible visual impairment in the elderly, affecting hundreds of millions of patients. Müller glia cells (MGC), the main type of glia found in the vertebrate retina, can resume proliferation in the adult injured retina and contribute to tissue repair. Also, MGC can be genetically reprogrammed through the expression of the transcription factor (TF) Achaete-scute homolog 1 (ASCL1) into induced neurons (iNs), displaying key hallmarks of photoreceptors, bipolar and amacrine cells, which may contribute to regenerate the damaged retina. Here, we show that the TF neurogenin 2 (NEUROG2) is also sufficient to lineage-reprogram MGC into iNs. The efficiency of MGC lineage conversion by NEUROG2 is similar to that observed after expression of ASCL1. However, reprogramming efficiency is affected by previous exposure to EGF and FGF2 during the expansion of MGC population. Transduction of either Neurog2 or Ascl1 led to the upregulation of key retina neuronal genes in MGC-derived iNs, but only NEUROG2 induced a consistent increase in the expression of putative retinal ganglion cell (RGC) genes. In vivo electroporation of Neurog2 in the neonatal retina also induced a shift in the generation of retinal cell subtypes, favoring the differentiation RGCs at the expense of MGCs. Altogether, our data indicate that Neurog2 induces lineage conversion of MGCs into RGC-like iNs.

## Introduction

The retina is a unique tissue with highly organized architecture, known to be one of the most energetically demanding systems in the nervous system (Wong-Riley, 2010). Due to oxidative stress, trauma or genetic mutations, gradual and irreversible cell death affects specific neuronal types in the retina (Athanasiou et al., 2013). Several neurodegenerative diseases affect the visual system and may lead to incurable blindness, such as *retinitis pigmentosa*, age-related macular degeneration and glaucoma (Bramall et al., 2010;Zhang et al., 2016). For instance, retinal ganglion cells (RGCs) degenerate in glaucoma eventually leading to blindness. In the last five years almost 65 million people worldwide were diagnosed with glaucoma (Gill et al., 2016;Liang et al., 2017), which is the leading cause of visual impairment in developed countries (WHO). Still, despite the social and economic burden of such diseases, therapeutic approaches are limited. Recent progress in cell-based therapy may, nonetheless, provide novel means to restore vision in glaucoma patients (Abu-Hassan et al., 2015;Chamling et al., 2016).

Cell lineage-reprogramming techniques, which allow the direct conversion of a non-neuronal cell into neurons, offer a powerful strategy to regenerate neuronal cells in the injured retina. In fact, expression of the bHLH neurogenic transcription factor (TF) Achaete-scute homolog 1 (ASCL1) *in vitro* induced the reprogramming of mouse Müller glia cells (MGC) into bipolar cells and, to a lesser extent, amacrine cells (Pollak et al., 2013). Following NMDA-mediated injury in postnatal mouse retina, ASCL1 expression reprogrammed MGCs into neurons expressing markers of bipolar cells, amacrine cells and photoreceptors (Ueki et al., 2015). Notably, when combined with the inhibitor of histone deacetylases trichostatin A, expression of ASCL1 elicited the conversion of some MGC into bipolar and amacrine cells in the injured adult retina (Jorstad et al., 2017). These findings demonstrate that regenerative effects of transgenic expression of ASCL1 in mouse Müller glia are akin to the regenerative response observed in non-mammalian vertebrates (Wilken and Reh, 2016). However, ASCL1 expression is not sufficient to reprogram MGCs into RGCs either in vitro or in vivo (9-11).

During development, expression of ASCL1 defines a subset of retinal progenitor cells (RPCs) that generate all neuronal types in the retina, except RGCs (Brzezinski et al., 2011). In contrast, expression of the bHLH TF Neurogenin 2 (NEUROG2) defines a separate set of RPCs, co-expressing the POU Class 4 Homeobox 1 and 2 (Pou4f1/Brn3a and Pou4f2/Brn3b) and contributing to the generation of RGCs (Hufnagel et al., 2010;Brzezinski et al., 2011). Interestingly, knocking down the expression of Pou4f2/Brn3b in MGCs cultured in conditions to induce stem cell-like properties hampers the differentiation into RGCs (Singhal et al., 2012;Song et al., 2013;Wu et al., 2016).

Here we report that forced expression of NEUROG2 is sufficient to convert MGC into a neurogenic state. Either ASCL1 or NEUROG2 elicited induced neurons (iNs) that express genes of bipolar, horizontal and amacrine cells, as well as photoreceptors. However, only forced expression of NEUROG2 led to the generation of iNs expressing hallmarks of RGCs. We also show that treatment with epidermal growth factor (EGF) and basic fibroblast growth factor (FGF-2) during the expansion of MGCs affects lineage-conversion efficiencies and iN-fate specification. Finally, we provide evidence for an instructive role of NEUROG2 in the specification of RGC fate *in vivo*. Collectively, our results may contribute to design new strategies to reprogram MGC into RGCs in the adult mammalian retina and, therefore, to the advance of gene-based therapies to treat retinal degenerations.

## Materials and Methods

### Animals

C57BL/6 mice were obtained from the Biotério Setorial do Instituto do Cérebro (BISIC). All experiments were approved by and carried out in accordance with the guidelines of the Institutional Animal Care and Use Committee of the Federal University of Rio Grande do Norte (license number #048/2014).

### Müller glial cell (MGC) culture

MGCs were purified from postnatal day (P)7-9 mice according to previously described protocols (de Melo Reis et al., 2008). Briefly, retinas were dissected out and chemically dissociated with TrypLE (Life Technologies) for 10 min at 37°C. Isolated cells were counted using a Neubeuer chamber and plated onto T75 culture flasks with DMEM F12 (Gibco) plus 10% fetal bovine serum (Gibco), 3.5 mM glucose (Sigma), 4.5g/L GlutaMax (Gibco), 100U/mL penicillin/streptomycin (Gibco), either with or without 10 ng/mL of epidermal growth factor (EGF, Gibco) and 10 ng/mL of fibroblast growth factor 2 (FGF2, Gibco). Half of the medium was changed once a week during the period of MGCs expansion.

### Plasmids

Plasmids contain the internal chicken β-actin promoter fused with a cytomegalovirus enhancer (pCAG), the coding sequence for either Ascl1 or Neurog2, an internal ribosomal entry site (I) and coding sequences for either DsRed or GFP (pCAG-Ascl1-I-DsRed, pCAG-Neurog2-I-DsRed and pCAG-Neurog2-I-GFP). Control plasmids encode only DsRed or GFP (pCAG-I-DsRed or pCAG-I-GFP).

Plasmid stocks were prepared in *Escherichia coli* and purified using an endotoxin-free Maxiprep plasmid kit (Invitrogen). DNA concentration was adjusted to 1 µg/µL in endotoxin free TE buffer, and plasmids were stored at −20°C.

### Nucleofection

After confluence, MGCs were chemically detached from T75 culture flasks with TrypLE enzyme at 37°C, and approximately 3×10^5^ cells were mixed with P3 solution (Lonza) and 1 µg of plasmids encoding for either NEUROGENIN2 (pCAG-Neurog2-IRES-DsRed) or ASCL1 (pCAG-Ascl1-IRES-DsRed) or only reporter protein DSRED (pCAG-IRES-DsRed). These solutions were placed in a special cuvette and electroporated using Nucleofector 4D (Lonza) with the P3 primary cell program. Next, 8 × 10^4^ cells were plated onto glass-coverslips (100013-Knittel) previously coated with laminin (L2020 - SIGMA) and Poly-D lysine (Sigma) containing pre-warmed DMEM F12, 10% fetal bovine serum, 3.5 mM glucose, 4.5g/L GlutaMax and 100U/mL penicillin/streptomycin. After 24h, medium was replaced with differentiation medium containing DMEM F12, 3.5 mM glucose, 4,5g/l GlutaMax, 100U/ml penicillin/streptomycin and 2% B27 (Gibco).

### In vivo electroporation

In vivo electroporation was performed as described previously (Matsuda and Cepko, 2004;2007). Briefly, P0 Lister hooded rats were anesthetized by placing on ice. One microliter of DNA mix (6.5 µg/µL) containing 0.1% Fast Green dye (Sigma) prepared in HBSS saline was injected into the subretinal space with a Hamilton syringe equipped with a 33G blunt end needle. Five 99 V pulses were administered for 50 ms at 950 ms intervals, using a forceps-type electrode (Nepagene, CUY650P7) with Neurgel (Spes Medica).

### Immunocytochemistry

Cell cultures were fixed in 4% paraformaldehyde (PFA) in PBS for 15 min at room temperature. The cells were incubated overnight at 4°C, with primary antibodies diluted in PBS, 0.5% Triton X-100 and 5% goat serum, washed and incubated for 2 h at room temperature, with species-specific secondary antibodies conjugated to Alexa fluorophores. To stain the nuclei, cells were incubated for 5 min with 0.1 µg/mL DAPI (4’6’-diamino-2-phenylindone) in PBS 0.1 M. Coverslips were mounted onto glass slides with Aqua Poly/Mount mounting medium (Polysciences, Warrington, PA). Primary antibodies and respective dilutions were: chicken anti-Green Fluorescent Protein (Aves Labs, cat#GFP-1020, 1:1000), rabbit anti-Red Fluorescent Protein (Rockland, cat#600- 401-379, 1:1000), mouse anti-microtubule associated protein (Sigma, cat#M1406, 1:500), mouse anti-β_III_-TUBULIN (TUBB3; Biolegend, cat#MMS-435P, 1:1000), rabbit anti-RBPMS (PhosphoSolutions, cat#1830; 1:100), mouse anti-SYNAPSIN 1 (Synaptic Systems, cat#106001, 1:2000), mouse anti-PARVALBUMIN (SIGMA, cat#p3088, 1:1000), mouse anti-CRALBP (ABCAM, cat# ab15051, 1:500), rabbit anti-GFAP (DakoCytomation, cat#z0334, 1:4000) rabbit anti-SOX2 (ABCAM, cat# ab97959, 1:500), rat anti-BrdU (ABCAM, cat# ab6326 1:500), rabbit anti-phospho-histone 3 (Millipore, cat#06-570, 1:1000), mouse anti-PAX6 (Millipore, cat#MAB5552, 1:500) and mouse anti-NESTIN (Millipore, cat#mab353, 1:200 millipore).

### Calcium imaging

Calcium imaging was done on MGC cultures at 2-3 weeks post-nucleofection, using Oregon green 488 BAPTA-1 (Invitrogen, 10µM). Imaging was performed in a physiological solution containing 140 mM NaCl, 5 mM KCl, 2 mM MgCl_2_, 2 mM CaCl_2_, 10 mM HEPES, 10 mM glucose, and 6 mM sucrose, and pH 7.35. Images were acquired every 10 ms with virtually no intervals using a scientific CMOS camera (Andor). The microscope was controlled by Micro-Manager software together with the image processor ImageJ. Changes in fluorescence were measured for individual cells using the time series analyzer plugin v3.0 in ImageJ v1.37. The average of the first ten time-lapse images for each region of interest (ROI) was defined as initial fluorescence (F_0_), and the variation of fluorescence (ΔF) in each frame (n) was calculated as F_n_-F_0_/F_0_.

### Quantitative RT-PCR

MGC cultures were harvested at 13 days after nucleofection, and mRNA was isolated from all cells, including non-transfected MGCs. RNA was extracted using RNeasy Mini Kit (QIAGEN, CA, USA), which includes a genomic DNA elimination step, and the purity and quantity of total RNA was estimated using a ND8000 spectrophotometer (Thermo Scientific NanoDrop Products, DE, EUA). Extractions were carried out of cells from each group (Control, Neurog2 or Ascl1), detached chemically with TrypLE and washed with nuclease free PBS, following the manufacturer’s protocol. The first-strand cDNA was synthetized using the High-Capacity cDNA Reverse Transcription Kit (Applied Biosystems, NY, USA) in accordance with the manufacturer’s instructions, using 900 ng of extracted RNA per sample. Conditions for each cycle of amplification were as follows: 10 min at 25°C; 120 min at 37°C, 5 min at 85°C. The final cDNA products were amplified using RT2 Real-Timer SyBR Green/ROX PCR Mix (QIAGEN, CA, USA) in 25 µL of a reaction mixture pipetted into each well of a 96-well in a Mouse RT2 Profiler Custom PCR Array. The array was designed to simultaneously examine mRNA levels of 18 genes commonly expressed in retina cell types (RLBP1, GLUL, NRL, RHO, RCVRN, PDE6G, PROX1, LHX1, VSX2, SLC32A1, TH, CHAT, SLC17A6, POU4F1, CALB2, RBFOX3, SYN1 and PVALB) and 2 housekeeping genes (GAPDH and RPL19), following the manufacturer’s protocol. Real-time PCR was performed using a two-step cycling program, with an initial single cycle of 95° C for 10 min, followed by 40 cycles of 95° C for 15 seconds, then 60° C for 1 min, in an ABI ViiA 7 Real-Time PCR System (Applied Biosystems, NY, USA) with Sequence Detector System software v1.2. The ramp rate was adjusted to 1°C/s following manufacturer’s instruction. A first derivative dissociation curve was built (95°C for 1 min, 65°C for 2 min, then ramped from 65°C to 95°C at a rate of 2°C/min). The formation of a single peak at temperatures higher than 80°C confirmed the presence of a single PCR product in the reaction mixture.

For data analysis the 2^(−ΔΔCt)^ method (Livak and Schmittgen, 2001) was implemented using normalized threshold cycle (Ct) values provided by two independent experiments of nucleofection. Furthermore, we applied experiments using mRNA pool of two independent transfection experiments and used the Ct data to perform normalization and follow the 2^(−ΔΔCt)^ method (Livak and Schmittgen, 2001), considering the sensitivity, specificity, and reproducibility expected of real-time PCR using RT^2^ Profiler PCR Array System from QIAGEN. Endogenous gene control used in the normalization was the average of the mouse GAPDH and RPL19. A positive value indicates gene up-regulation and a negative value indicates gene down-regulation.

To confirm that genes upregulated in MGCs could indicate the acquisition of a retinal neuron-like phenotype in MGC-derived iNs, we performed the same analysis using cerebellum astroglia cell cultures 13 days after nucleofection with either Neurog2 or Ascl1 (Chouchane et al., 2017).

### Bioinformatics

Dataset raw count table and published metadata were obtained from GSE63472 accession code. A modified Seurat pipeline was used to re-analyze single-cell RNAseq data. First, we excluded from our analysis genes that were not expressed in at least 10 cells. Next, we selected cells expressing 500 to 5000 genes, and less than 5% of mitochondrial genes (n = 21494 cells). Metadata variables as number of genes and percentage of mitochondrial expression were also used to regress out some unnecessary clustering bias. Based on *PCElbowPlot*, we used 30 PC’s in *FindClusters* (resolution = 2) and *RunTSNE* Seurat’s functions. After that, using old assigned clusters and markers found by *FindAllMarkers* function (Macosko et al., 2015), new assigned clusters were labeled. Retinal cell types (n = 21176 cells) were classified according to the levels of expression of genes in Müller glia cells, astrocytes, amacrine cells, bipolar cells, horizontal cells, cones, rod cells and ganglion cells (Macosko et al., 2015). Cones and rod cells were merged in a single-group (Photoreceptors).

### Quantifications

To characterize MGC cultures we examined 20 fields at 40x magnification for Cralbp and GFAP, and 20 fields at 20x magnification for Nestin, Pax6 and PH3 proteins. To quantify the reprogramming process, cells were examined for colocalization of DSRED and TUBB3 immunoreactivity at 13 days post nucleofection (dpn), in 20 fields at 20x magnification, and the same was done for MAP2 protein. For all protocols of quantification, we counted immunoreactive cells in 3 independent experiments.

To estimate a neuronal polarization index, we divided the neurons into 4 quadrants and measured the axial distribution of neuronal processes. Nineteen neurons were analyzed for the condition MGC with EGF/FGF + Neurog2, 31 neurons for MGC without + Neurog2, 26 neurons for MGC with EGF/FGF + Ascl1 and 21 neurons for MGC without EGF/FGF + Ascl1.

Distribution of GFP+ cells in the P10 rat retinas following in vivo electroporation at P0 in Lister-hooded pups was examined in 26 sections from 5 control-electroporated retinas and 27 sections from 5 Neurog2-electroporated retinas. The outer nuclear layer (ONL), inner nuclear layer (INL) and ganglion cell layer (GCL) were identified using DAPI counter-staining.

### Statistical analysis

All statistical data are presented as the mean ± standard error of the mean (SEM) of at least three independent experiments. Statistically significant differences were assessed using unpaired t-test or one-way or two-way analysis of variance (ANOVA). Confidence interval is 95%.

## Results

### Properties of cells expanded in the presence or absence of EGF/FGF2

Enrichment of MGCs was done according to a previously described method (Hicks and Courtois, 1990). Using this method we (de Melo Reis et al., 2008) and others (Das et al., 2006) have shown that virtually all cells after 2-3 weeks in culture express the MGC markers vimentin, glutamine synthetase (GS), cellular retinaldehyde-binding protein (CRALBP). Based on the previously reported use of selected growth factors to expand astroglial populations for lineage reprogramming into iNs (Berninger et al., 2007;Heinrich et al., 2010;Chouchane et al., 2017), we cultured cells either with or without EGF/FGF2. After expansion, virtually all cells in cultures obtained using both protocols expressed the MGC protein CRALBP (Fig 1A,B). A high percentage of cells also expressed glial acid fibrillary protein (GFAP), which is upregulated in MGC cultures (Das et al., 2006). No significant difference was observed between the two expansion conditions (Fig 1A,B).

**Fig 1.**
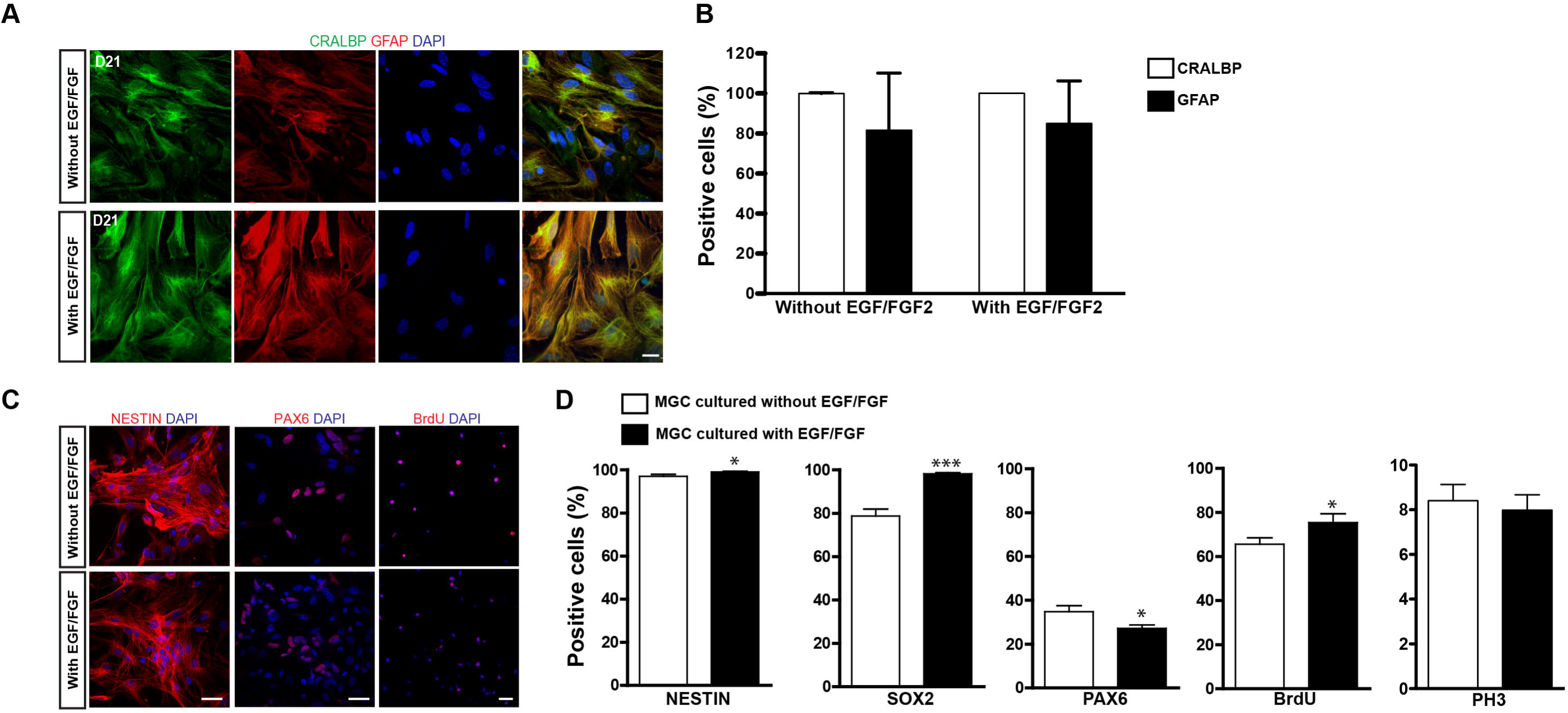
Characterization of MGC cultures expanded with or without EGF/FGF2. (A) Representative photomicrographs of MGC cultures showing the expression of CRALBP (green) and GFAP (red) 21 days after expansion in the absence (top) or presence (bottom) of EGF/FGF2. (B) Frequencies of CRALBP and GFAP positive cells in enriched MGC cultures. (C) Immunolabeling for NESTIN, PAX6 and BRDU (red) in MGCs expanded in the absence (top) or presence (bottom) of EGF/FGF2. (D) Frequencies of MGCs immunolabeled for NESTIN, SOX2, PAX6, BrdU and PH3 prior to lineage reprogramming (**p<0.01; ***p<0.001; Unpaired t-test). Scale bars: 20µm

MGCs and late-progenitors in the retina share the expression of several proteins, including NESTIN, sex determining region Y (SRY)-box 2 (SOX2) and Paired box 6 (PAX6) (Karl et al., 2008;Lin et al., 2009;Bhatia et al., 2011), what has led to the suggestion that MGCs may have neural stem cell-like properties (Das et al., 2006;Lawrence et al., 2007;Nickerson et al., 2008;Giannelli et al., 2011). Accordingly, we found that almost all cells expanded in our culture systems expressed NESTIN (Fig 1C) and SOX2, but the proportion of cells expressing these proteins was slightly higher in cultures grown in the presence, rather than in the absence of EGF/FGF2 (Fig 1D). In contrast, only a fraction of MGCs expressed the PAX6 protein, and this fraction was higher in the absence of EGF/FGF2 (Fig 1C, D).

To examine the proliferative potential of enriched MGCs, we added the thymidine analog BrdU to the cultures at day 20. The percentage of MGCs labeled with BrdU (BrdU+) at 24 h of incubation was slightly higher with, rather than without EGF/FGF2 (Fig 1C,D). However, the percentage of phospho-histone labeled (PH3+) mitotic MGCs was similar in both conditions (Fig 1D), what could be explained by a selective lengthening of the S-phase upon EGF/FGF2 treatment. Collectively, these observations support the interpretation that cells enriched in our cultures are presumptive MGCs that retain some properties observed in late-progenitors of the developing retina. However, we never observed spontaneous neurogenesis in these cultures, suggesting that a potential progenitor state of presumptive MGCs in culture is associated with a glial-fate restriction.

### Expression of either NEUROG2 or ASCL1 is sufficient to convert MGCs into iNs

Next, we tested whether the expression of NEUROG2 may reprogram enriched MGCs into iNs, as compared with ASCL1 as previously described (Pollak et al., 2013). Expanded MGCs were harvested and transfected with control-I-GFP, Neurog2-I-GFP (Fig 2B), Neurog2-I-DsRed (Figs 2D and F) or Ascl1-I-DsRed plasmids (Fig 3A and C), and maintained for 13 days (1 day in growth medium + 12 days in differentiation medium; Fig 2A). At the end of this period, cells transfected with control plasmids maintained both their typical glial morphology and the content of the glia-specific protein GFAP (Fig 2C). In contrast, a substantial fraction of NEUROG2-containing cells underwent robust morphological changes, characterized by reduction of the cell body and extension of thin and long primary processes (usually 2 or 3) with small ramifications, similar to the morphology of neuronal cells in culture (Figs 2B, D and F). Accordingly, these cells also contained the neuron-specific proteins β_III_-TUBULIN (Fig 2D) and microtubule associated protein 2 (MAP2) (Fig 2F), indicating that NEUROG2 led to conversion of cultured MGCs into iNs. The frequency of iNs was significantly higher in cultures containing MGCs expanded in the presence, as compared with the absence of EGF and FGF2 (71.0 ± 4.1% versus 26.4 ± 5.1%; p<0.001; Tukey’s multiple comparison test; Fig 2E), suggesting that these growth factors facilitate reprogramming. Additionally, the vast majority of iNs also expressed MAP2 (Fig 2F and G) independently of mitogenic factors during enrichment.

**Fig 2.**
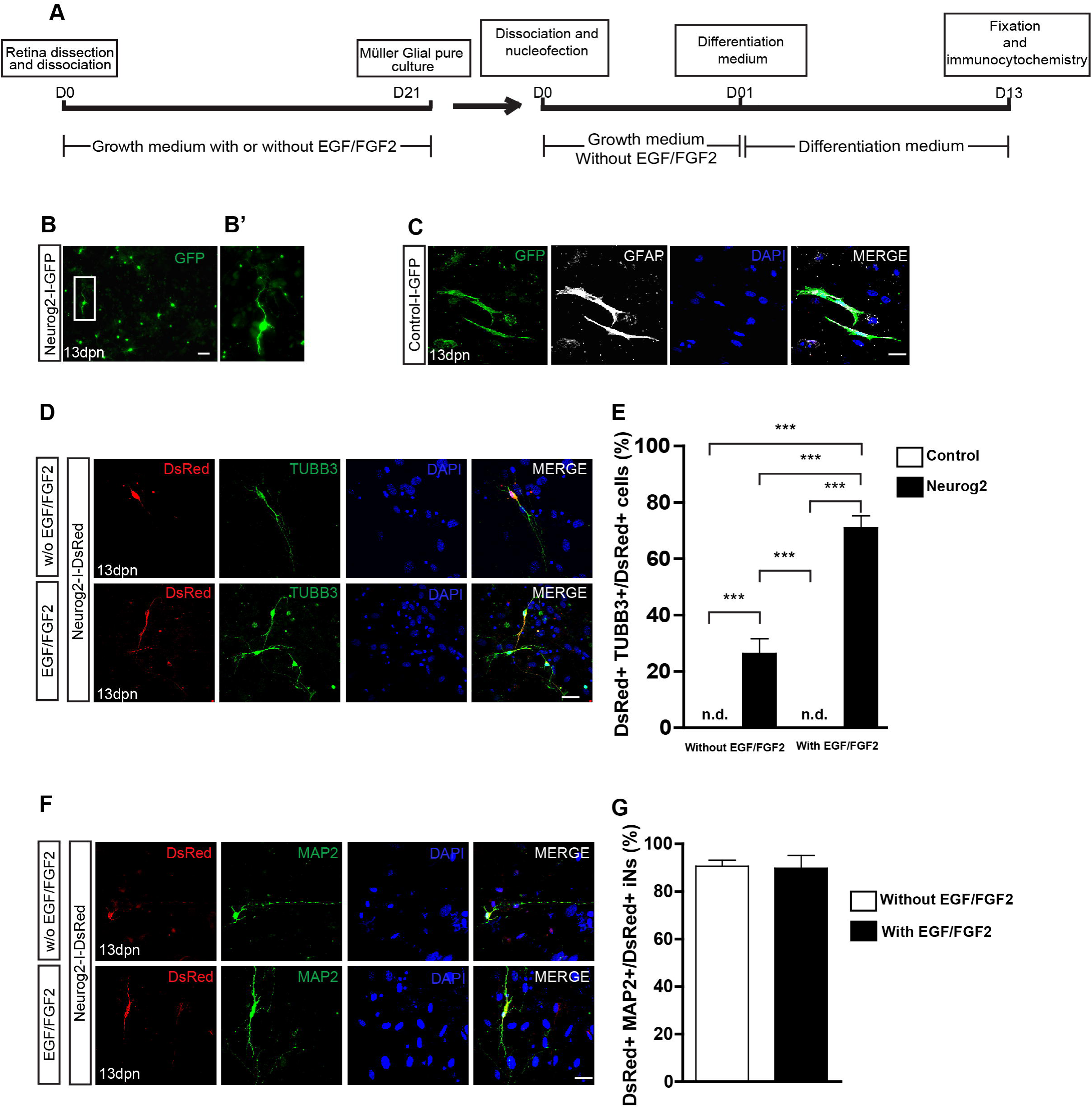
Expression of NEUROG2 converts MGCs into iNs. (A) Experimental design. Note that EGF/FGF2 are used only during MGC expansion. (B-B’) Overview of MGC cultures 13 days post nucleofection (dpn) with Neurog2-I-GFP showing the presence of GFP+ cells (green) adopting a neuronal-like morphology (white inset, magnified in B′). (C) Representative photomicrographs of MGC cultures at 13 dpn with control-I-GFP, showing GFP+ cells (green) that maintain glial morphology and contain GFAP (white). (D) Immunolabeling of the neuronal protein β_III_-TUBULIN (TUBB3, green) in DsRed (red) cells at 13 dpn with Neurog2-I-DsRed in MGCs cultures expanded either without (top) or with (bottom) EGF/FGF2. (E) Frequencies of DsRed+/TUBB3+ iNs amongst total DsRed+ cells at 13 dpn (***p<0.001; Tukey’s multiple comparison test). (F) Immunolableing for the neuronal protein MAP2 (green) in DsRed (red) cells at 13 dpn with Neurog2-I-DsRed in MGCs expanded either without or with EGF/FGF2. (G) Frequencies of DsRed+MAP2+ iNs amongst all DsRed+ iNs. Photomicrographs in C, D and F are single confocal Z stacks. Nuclei are stained with DAPI (blue). Scale bars: 20µm.

**Fig 3.**
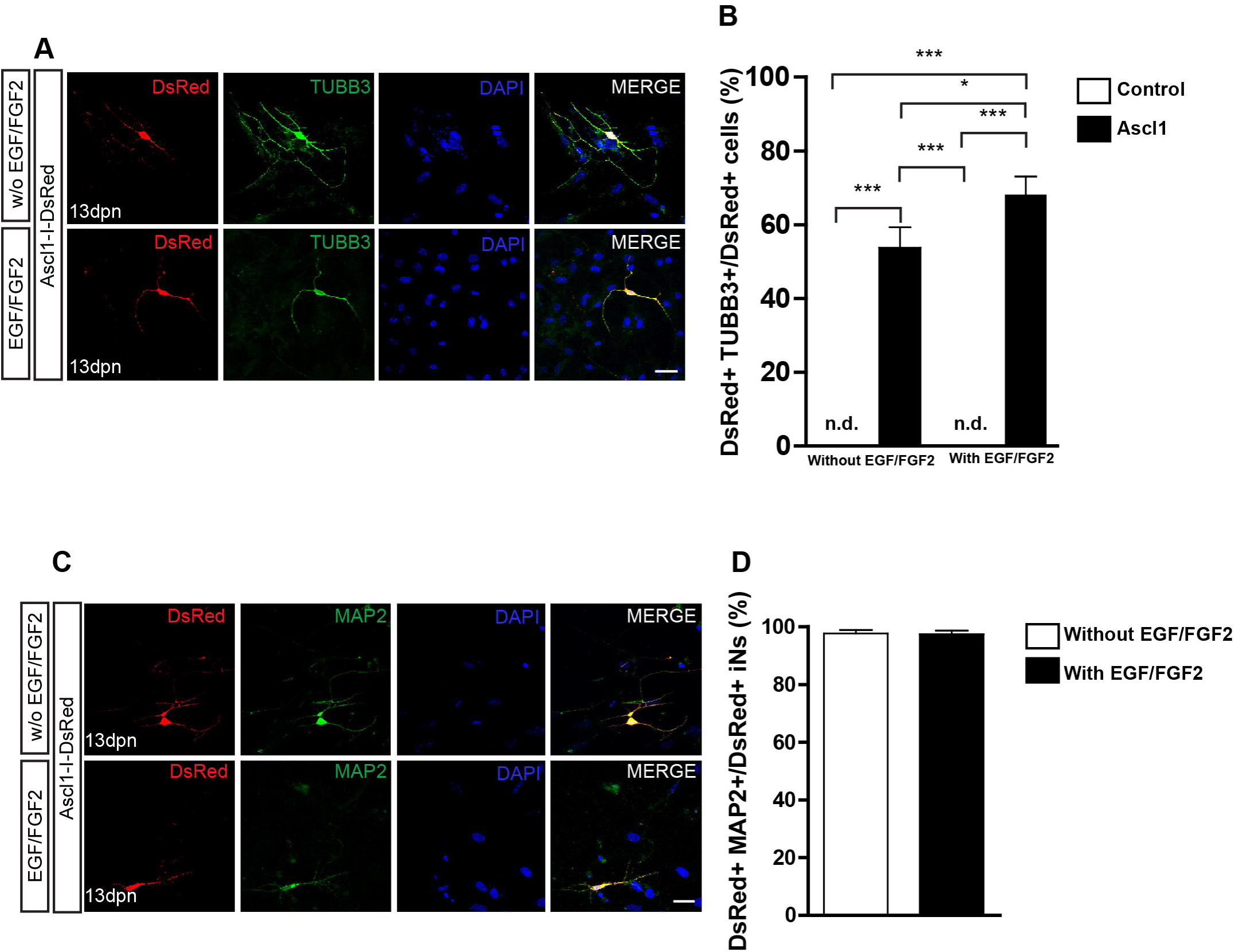
Expression of ASCL1 converts MGCs into iNs. (A) Representative photomicrographs of MGC cultures expanded without or with EGF/FGF2 at 13 dpn with Ascl1-I-DsRed, showing DsRed+ cells (red) immunolabeled for β_III_-TUBULIN (TUBB3, green) (B) Frequencies of DsRed+/TUBB3+ iNs amongst total DsRed+ cells at 13 dpn (*p<0.05; ***p<0.001; Tukey’s multiple comparison test). (C) Immunolabeling of the neuronal protein MAP2 (green) in DsRed (red) cells at 13 dpn with Ascl1-I-DsRed in MGCs expanded without or with EGF/FGF2. (G) Frequencies of DsRed+MAP2+ iNs amongst all DsRed+ iNs. Photomicrographs in A and C are single confocal Z stacks. Nuclei are stained with DAPI (blue). Scale bars: 20µm.

Lentiviral-mediated expression of ASCL1 reportedly induced the conversion of about 30% of cultured MGCs into neuronal-like cells (Pollak et al., 2013). In our model, nucleofection of ASCL1 led more than half of cultured MGCs to convert into iNs (Fig 3 A,B). The proportion of iNs was higher among MGCs previously expanded in the presence, rather than in the absence of EGF and FGF2 (68.1 ± 4.9% versus 53.7 ± 5.5%; p<0.05; Tukey’s multiple comparison test; Fig 3B), and virtually all iNs also expressed MAP2 (Fig 3D). Collectively, our data indicate both that the expression of either NEUROG2 or ASCL1 is sufficient to lineage-reprogram MGCs into iNs, as well as that previous exposure of MGCs to EGF and FGF2 facilitates reprogramming.

### Functional and morphological differentiation of MGC-derived iNs

To evaluate whether iNs derived from lineage reprogrammed cultured MGCs develop features of mature functional neurons, we performed calcium imaging using a genetically encoded calcium indicator (GCAMP5) and a fluorescent dye (Oregon green BAPTA-1). MGC cultures nucleofected with either Ascl1 or Neurog2 expression plasmids were maintained for 15 days, and then treated with Oregon green BAPTA (see Materials and Methods). Fast fluctuations of intracellular calcium levels leading to sudden fluorescence changes, likely produced by an abrupt aperture of voltage-gated calcium channels mediated by synaptic activity (Bonifazi et al., 2009;Yang and Yuste, 2017), were detected in more than half of iNs (Neurog2 with EGF/FGF2: 12/20; Neurog2 without EGF/FGF2: 13/18; and Ascl1 with EGF/FGF2: 11/20; Ascl1 without EGF/FGF: 12/19) (Fig 4 and Supplementary Movies). In contrast, non-transfected MGCs showed slow calcium fluctuations (Fig 4).

**Fig 4.**
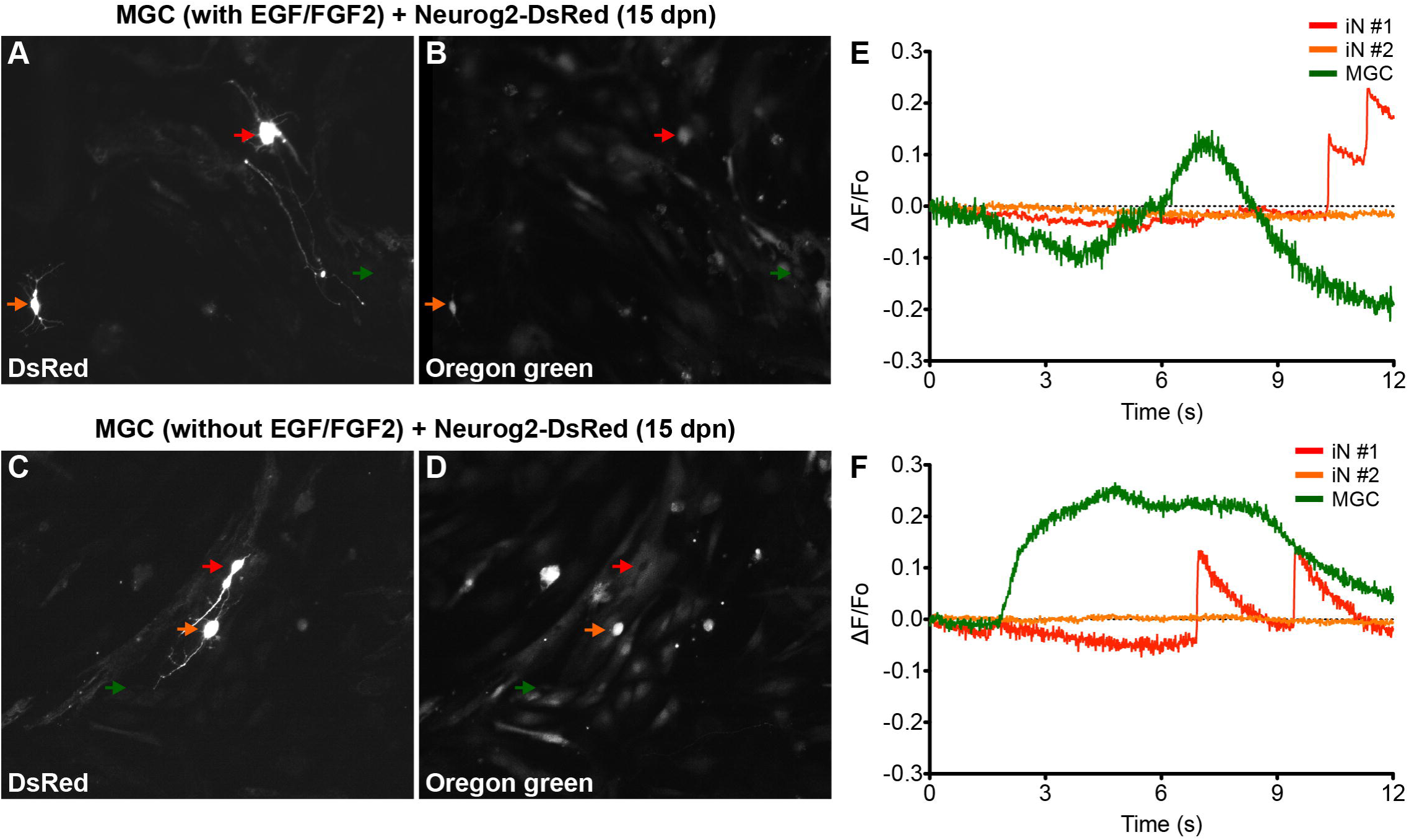
MGC-derived iNs display fast calcium transients. (A-D) Photomicrographs show examples of MGC-derived iNs at 13 days dpn with Neurog2-I-DsRed expressing DsRed (A and C), and labeled with Oregon Green Bapta (B and D). Red and orange arrows point to DsRed+ MGC-derived iNs, whereas green arrows point to non-transfected MGCs randomly selected within the same fields of observation. Top (A-B) and bottom (C-D) images were obtained from MGCs expanded in the presence of absence of EGF/FGF2, respectively. (E-F) Representative traces of calcium-transients in cells indicated by colored arrows in panels A (E) and C (F). Calcium responses were calculated as the change in fluorescence (ΔF) over the initial fluorescence (F_0_). Note that only iNs indicated by the red arrows displayed fast calcium-transients as indicated by rapid variations in the fluorescence (raising interval between 10-30 ms). In contrast, non-transfected MGCs (green arrows) have only slow calcium-transients as indicated by gradual changes in fluorescence levels (raising interval > 1s), suggestive of metabotropic responses. DsRed+ cells indicated by the orange arrows did not show any significant fluctuation in fluorescence levels during the period of observation.

We also examined the morphology of MGC-derived iNs in the various experimental groups (Fig S1). Expression of either Neurog2 or Ascl1 leads to the generation of diverse iNs (Figs S1A,B). Notwithstanding, whereas three-quarters of Neurog2-converted iNs were either unipolar or bipolar, Ascl1-converted iNs had approximately equal proportions of either multipolar or uni/bipolar morphologies (Fig S1C). These observations suggest that Neurog2 and Ascl1 iNs may acquire distinct phenotypes.

### MGC-derived iNs express key genes of retinal neurons

It has been suggested that the origin of reprogrammed astroglial cell lineages affects the phenotype of induced neurons (Chouchane et al., 2017). We tested whether MGC-derived iNs show hallmarks of bona fide retina neurons, after lineage conversion induced by either ASCL1 or NEUROG2 (Fig 5). First, we took advantage of single-cell transcriptomes available in the literature (Macosko et al., 2015) to identify molecular signatures of the major retinal cell types (Fig 5A and Fig S2). Based on single-cell RNA sequence of adult mouse retina cells (Macosko et al., 2015), we defined a small set of transcripts with enriched expression in MGCs (RLBP1 and GLUL), astrocytes (GFAP), photoreceptors (NRL, RHO, RCVRN and PDE6G), horizontal cells (PROX1 and LHX1), bipolar cells (VSX2), amacrine cells (PAX6, SLC32A1, GAD1, GAD2, TH and CHAT) and RGCs (RBPMS, SLC17A6, POU4F1, CALB2, RBFOX3, SYN1, TUBB3 and PVALB). Next, we used qRT-PCR to compare the expression of these molecular markers in MGC cultures expanded in the presence or absence of EGF/FGF2, and nucleofected with either Neurog2 or Ascl1 (Fig 5B and C). Despite the small number of iNs amongst the total population of MGCs (∼1-2%), we detected an increase in the expression of several genes commonly expressed by retina neurons in MGC cultures nucleofected with Neurog2 or Ascl1 (Figs 5B and C), but not among cerebellum astroglial cells nucleofected with the same plasmids (Fig S3), which are known to adopt mostly a GABAergic iN phenotype (Chouchane et al., 2017). Interestingly, differences were observed among cells converted through each of the two expansion protocols. While MGC-derived iNs, induced by expression of either ASCL1 and NEUROG2 grown in the absence of EGF/FGF2 upregulated key genes of photoreceptors, genes of amacrine cells were expressed only in iNs derived from MGCs expanded in the presence of growth factors. In contrast, increased levels of genes commonly expressed by horizontal and bipolar cells were detected in all experimental conditions. Surprisingly, we also found that many genes commonly expressed by RGCs were upregulated in MGC-derived iNs, in particular the specific RGC genes SLC17A6 and POU4F1 (Quina et al., 2005;Martersteck et al., 2017), but the same was not observed in cerebellar astroglia-derived iNs (Fig S3). This effect was much more robust after expression of NEUROG2 (Fig 5B), independently of the expansion protocol. Altogether, the transcriptome panel suggests that either Neurog2 or Ascl1 may induce distinct, but partly overlapping retina neuronal phenotypes in MGC-derived iNs, but Neurog2 seems to induce a more complete RGC-like phenotype.

**Fig 5.**
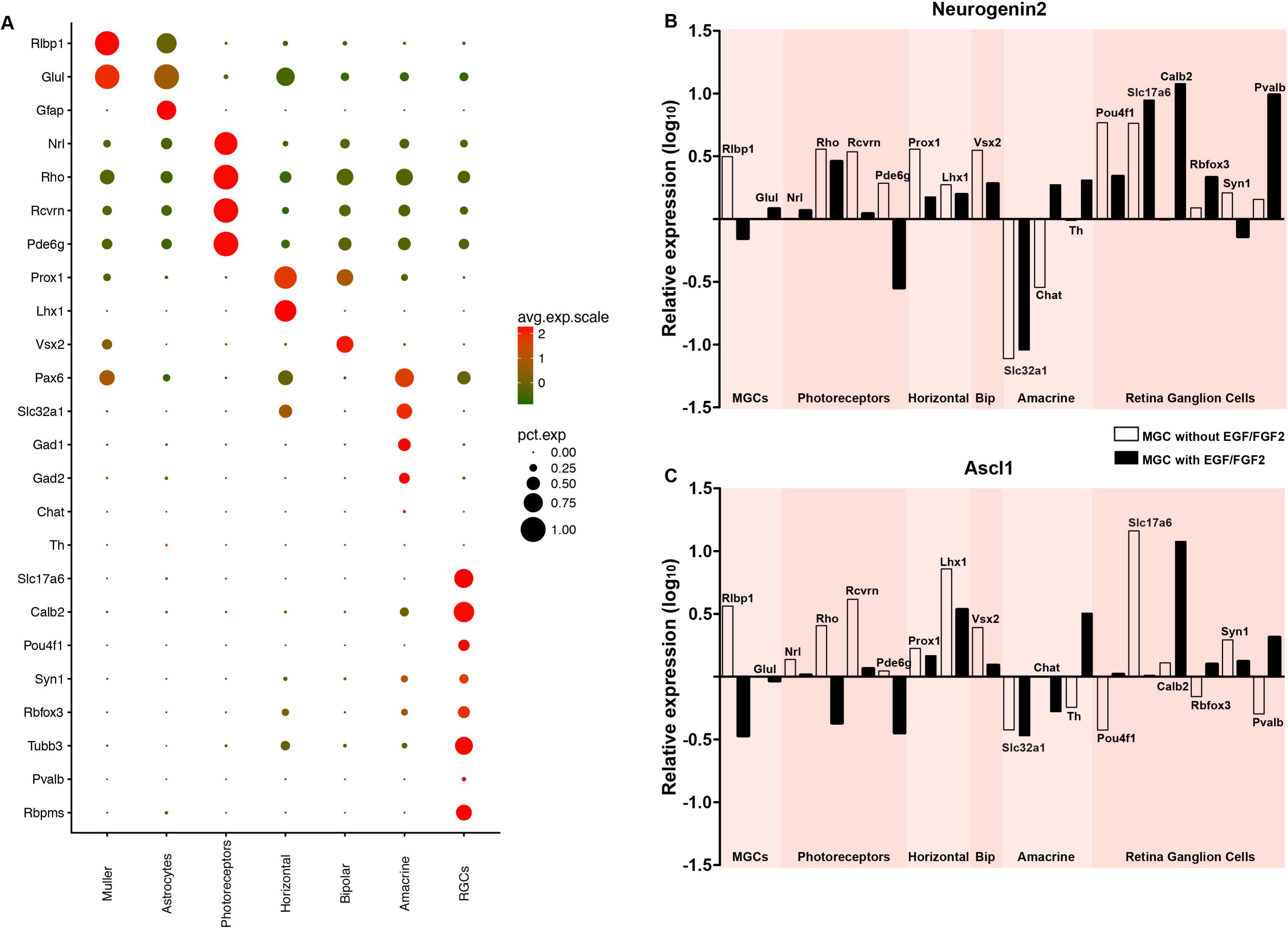
Expression of either NEUROG2 or ASCL1 induces upregulation of key retina neuron genes in MGC-derived iNs. (A) Expression levels of selected genes used to identify different classes of retina neurons using scRNAseq (see also Fig S4). Dot sizes and colors (green-low to red-high) represent the percentage of cells expressing the gene and its average scaled expression, respectively. Observe that the expression of Slc17a6 and Pou4f1 are specific for a high percentage of cells assigned as RGCs. (B-C) Graphics showing the relative expression levels (log_10_) of genes enriched in Müller glia (Rlbp1, Glul), photoreceptors (Rho, Rcvrn, Pde6g), horizontal (Prox1, Lhx1), bipolar (Vsx2), amacrine (Slc32a1, Chat, Th) and retina ganglion cells (Pou4f1, Slc17a6, Calb2, Rbfox3, Syn1, Pvalb, Rbpms), as shown in A, in MGC cultures nucleofected with either Neurog2 (B) or Ascl1 (C). White and black bars indicate that MGC were expanded in the absence or presence of EGF/FGF2, respectively, prior to the expression of Neurog2 or Ascl1.

### Neurog2-induced RGC-like phenotypes

To further test for the acquisition of RGC features by MGC-derived iNs, we immunolabeled markers of retina neurons in individual iNs (Fig 6). NEUROG2-converted neurons contained RNA Binding Protein mRNA Processing Factor (RBPMS; Figs 6A-C), a selective marker of RGCs in the mammalian retina (Rodriguez et al., 2014), as well as β_III_-TUBULIN (Figs 2 and 6D-F), PARV (Figs 6G-I) and SYN1 (Figs 6J-L), all of which are enriched in RGCs in vivo (Fig 5A). These findings, together with the qRT-PCR data, support the interpretation that NEUROG2 is sufficient to induce a RGC-like phenotype in MGC-derived iNs.

**Fig 6.**
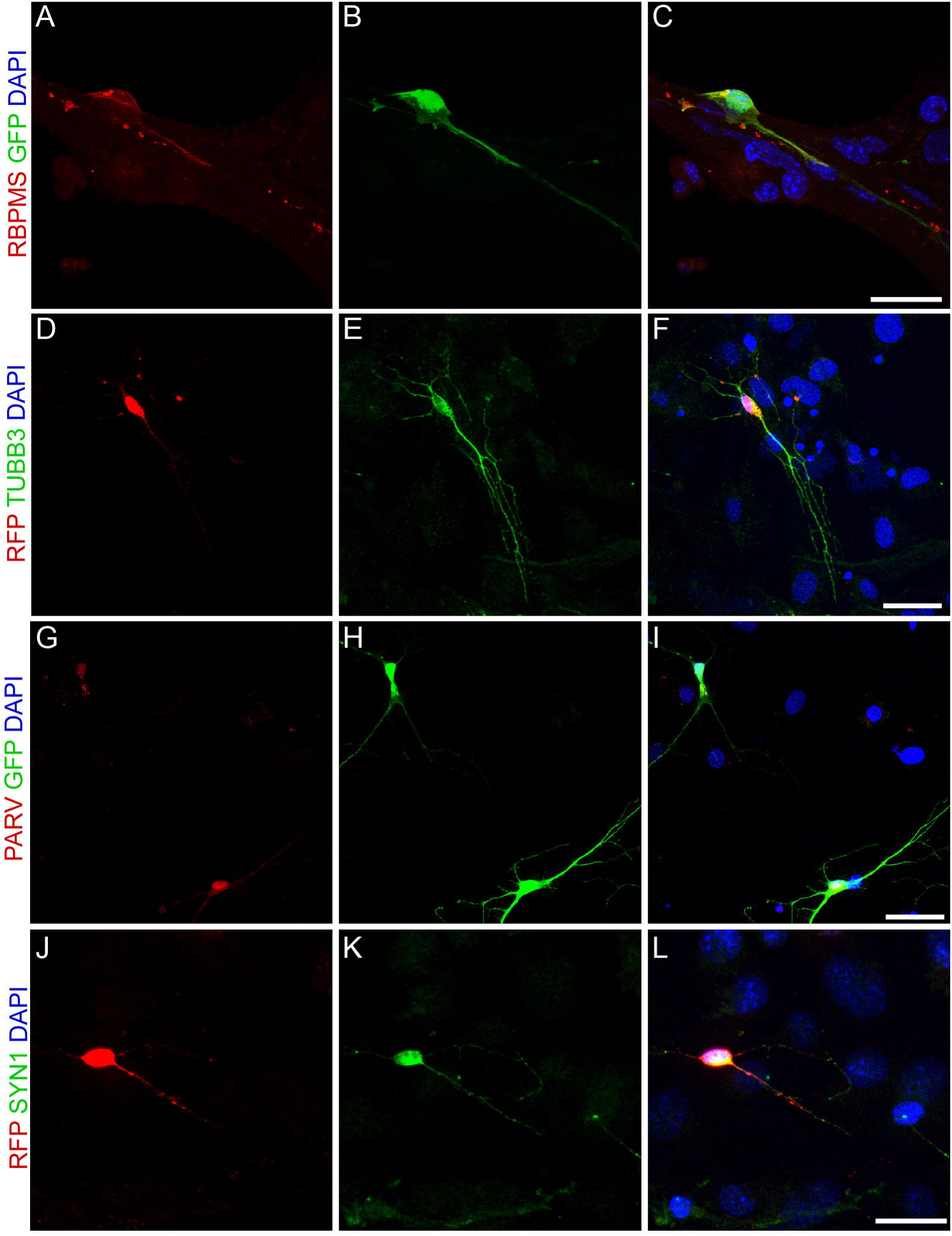
Expression of NEUROG2 induces the acquisition of RGC-like phenotypes in MGC-derived iNs. (A-L) Immunolabeling of the RGC proteins RBPMS (red, A-C), β_III_-TUBULIN (green, D-F), PARV (red, G-I) and SYN1 (green, J-L) in MGC converted into iNs at 15 days post-nucleofection with either Neurog2-I-GFP (green, A-C and G-I) or Neurog2-I-DsRed (red, D-F and J-L). Scale bars: 20 µm.

### Overexpression of NEUROG2 resumes RGC generation in the neonatal retina ***in vivo***

Finally, we tested whether the expression of NEUROG2 in late retinal progenitors induces *de novo* generation of RGCs. To that effect, sub-retinal injections of either pCAG-I-GFP or pCAG-Neurog2-I-GFP plasmids were followed by electroporation in P0 rats (Fig 7). As previously described (Matsuda and Cepko, 2004), the vast majority (∼80%) of GFP+ cells in control-electroporated retinas settled in the outer nuclear layer (ONL) and differentiated into rod cells, while the remaining cells located mostly in the inner nuclear layer (INL) and differentiated into bipolar, MGCs or amacrine cells (Figs 7A and E). Also consistent with the literature (Matsuda and Cepko, 2004), a very low number of GFP+ cells was found in the ganglion cell layer (GCL) of control-electroporated animals (2 cells in one animals out of 1254 GFP+ cells counted from 5 animals − 26 × 250 µm sections), whereas a significant proportion of cells in all animals displayed a morphology suggestive of MGCs (8 ± 3 %, n=5 animals, 1254 GFP+ cells). In contrast, in retinas electroporated with pCAG-Neurog2-I-GFP plasmid, we found a higher number of GFP+ cells in the GCL (17 cells in 5 animals out of 1340 GFP+ cells counted from 5 animals − 27 × 250 µm sections), and a complete absence of GFP+ MGCs (Fig 7C-I, Figs S4 and S5). The mean number of GFP+ cells within the GCL was significantly increased in Neurog2-electroporated animals as compared to controls, suggesting that NEUROG2 expression resumed the generation of RGCs (Fig 7F). Accordingly, we also found that GFP+ cells within the GCL expressed classical markers of RGCs, such as RBPMS (Fig 7 G-I) and β_III_-TUBULIN (Fig S4) and extended thin axonal processes that reached the optic nerve (Fig 7J). Some GFP-expressing cells within the GCL also extended processes towards the inner plexiform layer, consistent with the morphology of RGCs (Fig 7G and Fig 8). Moreover, GFP+ cells within the GCL also displayed varied patterns of dendrite stratification (Fig 8), consistent with distinct RGC morphologies in the rodent retina (Sanes and Masland, 2015). Altogether, these data support the hypothesis that the expression of NEUROG2 endows late retinal progenitors with the potential to generate RGCs, while it inhibits the differentiation of MGCs *in vivo*.

**Fig 7.**
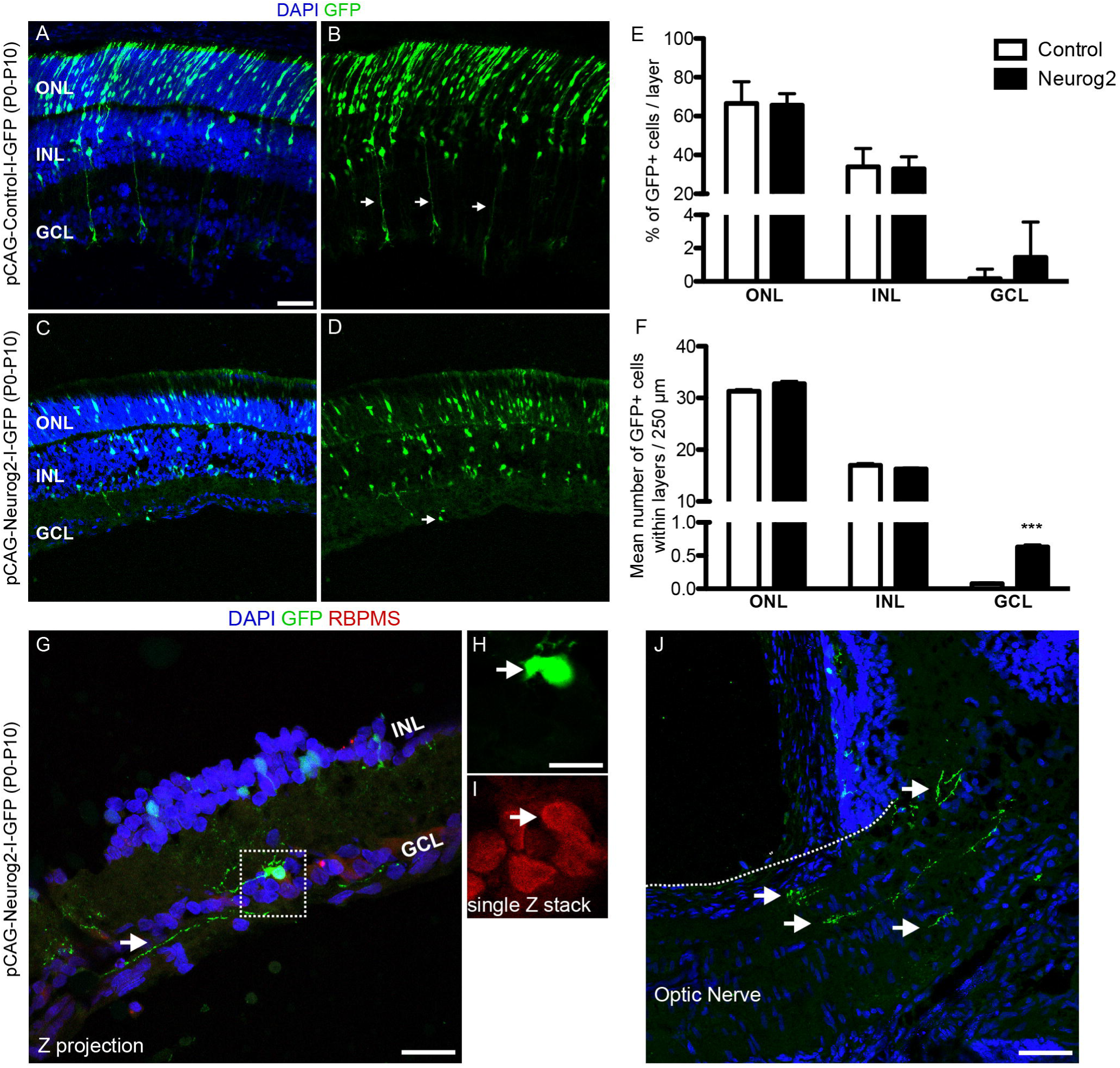
Forced-expression of NEUROG2 resumes the generation of RGCs in the developing retina *in vivo*. (A-D) Representative images of P10 rat retinas after electroporation with control-I-GFP (A,B) and Neurog2-I-GFP (C,D) plasmids at P0. Note the presence of GFP+ radial fibers typical of MGCs (arrows in B) and the complete absence of GFP+ cells in the GCL in controls. Observe also the decrease of GFP+ cells in the ONL and the presence of GFP+ cells in the GCL of Neurog2-electroporated retina (arrow in D). (E) Frequencies of GFP+ cells in the ONL, INL e GCL of the retina of control- and Neurog2-electroporated animals. (F) Mean numbers of GFP+ cells within the ONL, INL or GCL per 250 µm longitudinal section (***p<0.001; Two-way ANOVA, Bonferroni post-hoc test). (G-I) Expression of RBPMS (red) in GFP+ cells (green) within the GCL of Neurog2-electroporated retinas. White arrow points to the axon-like process of the GFP+ cell observed within the GCL. Dashed box (G) is amplified in H and I to show the co-localization of GFP and RBPMS in a single confocal Z-stack. (J) Axon-like GFP+ processes (arrows) reaching the optic nerve in a Neurog2-electroporated retina. Nuclei are stained with DAPI (blue). Scale bars: A-D and J: 50 µm; G: 25 µm; H-I: 12,5 µm.

**Fig 8.**
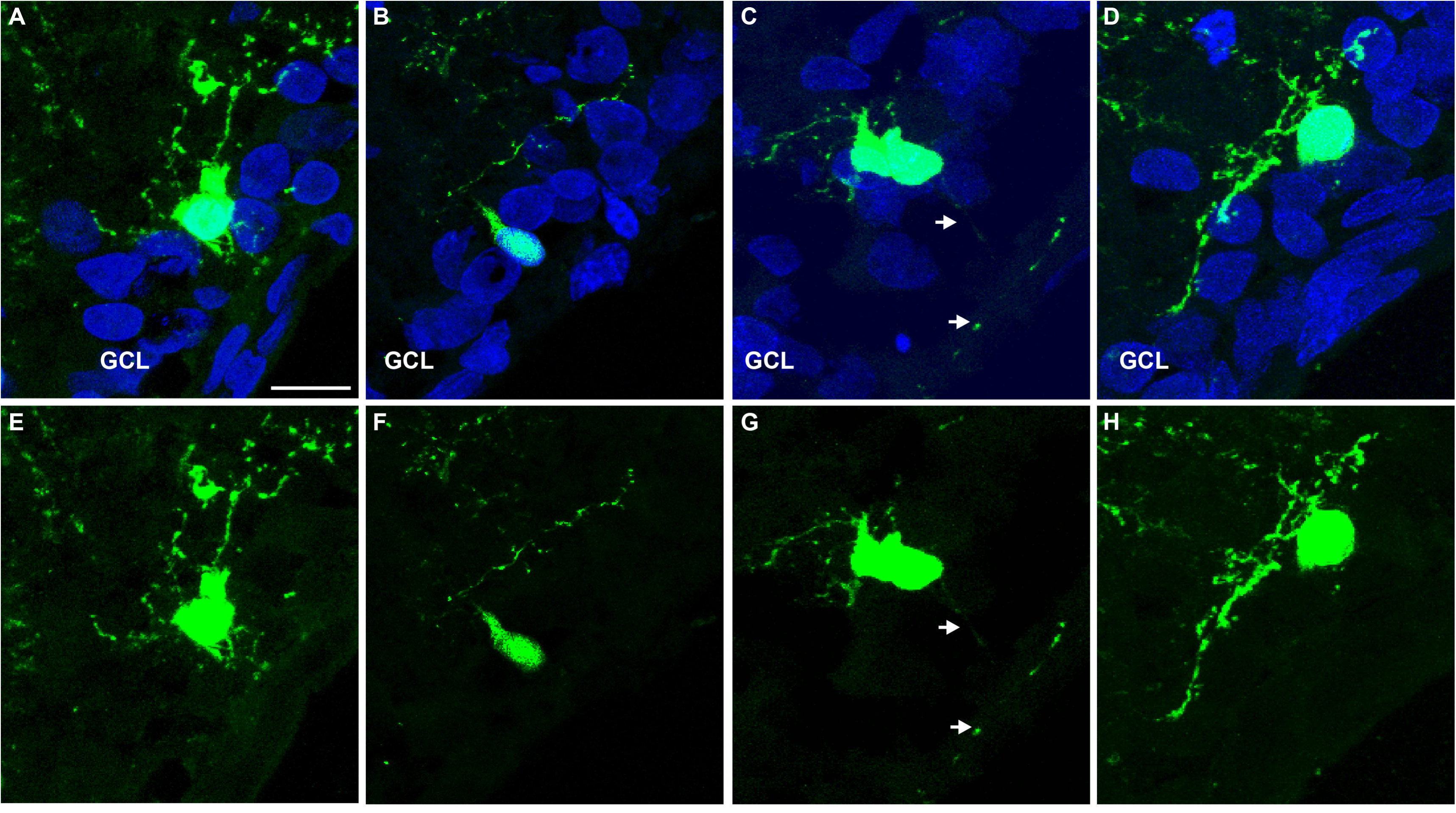
Representative examples of GFP+ cells within the GCL of Neurog2-electroporated retinas. (A-H) Images show GFP+ cells (green) within the GCL of P10 rats after electroporation with Neurog2-I-GFP at P0. Nuclei are stained with DAPI (blue, A-D). Note the distinct positions and dendrite stratifications of GFP+ cells (E-H), resembling *bona fide* subtypes of rodent RGCs (Sanes and Masland, 2015). Scale bar: 25 µm.

## Discussion

Here we show that cells isolated in culture and expressing several hallmarks of Müller cells, the main type of glia found in the vertebrate retina, can be reprogrammed into neurons when transduced with plasmids encoding either of the basic Helix loop Helix (bHLH) transcription factors Neurog2 or Ascl1. Cell–lineage reprogramming is affected by treatment with EGF and FGF2 during MGC enrichment, and led to the expression of typical neuronal markers, MAP-2 and β_III_-TUBULIN. Induced neurons expressed clusters of genes consistent with the profile of retinal cells, suggesting that distinct retinal neuron phenotypes are elicited in MGC-derived iNs. Notably, only NEUROG2-iNs upregulated a set of genes compatible with RGC identity. A role for NEUROG2 in the induction of RGC fate was also observed in the neonatal mouse retina *in vivo*, suggesting that Neurog2 is a good candidate for gene therapies aiming at the regeneration of RGCs in the adult injured retina.

Lineage reprogramming of astroglial cells into neurons has been heralded as a possible treatment for degenerative or traumatic brain disorders, and recent studies showed evidence of conversion of glial cells into functional neurons with high efficiency in the murine brain, induced by virally delivered transcription factors (Torper and Gotz, 2017). Astrocytes obtained from the postnatal rodent cerebral cortex (Berninger et al., 2007;Heinrich et al., 2010) and cerebellum (Chouchane et al., 2017), as well as retinal MGCs (Singhal et al., 2012;Pollak et al., 2013;Song et al., 2013;Ueki et al., 2015;Wu et al., 2016;Jorstad et al., 2017) have been reprogrammed into neurons.

Lentiviral gene transfer of the proneural factor Ascl1 partially reprogrammed P11/12 Müller glial cells *in vitro* into retinal progenitors 3 weeks after infection (Pollak et al., 2013), a process facilitated by microRNA miR-124-9-9* (Wohl and Reh, 2016). Combination of Ascl1 and miR-124-9-9* led to a peak of 50-60% reprogrammed iNs, whereas Ascl1 alone reached 30-35% (Wohl and Reh, 2016). The latter is lower than what we found following expression of Ascl1, which may be due to the earlier age of the mice from which cells were isolated in our study (P7-9). Indeed, in our hands, MGCs isolated from P21 retina failed to reprogram into iNs upon expression of either ASCL1 or NEUROG2 (data not shown), similar to what has been described for astroglia isolated from the adult mouse cerebral cortex (Heinrich et al., 2010) or even for astroglia isolated from the postnatal cerebral cortex or cerebellum, and maintained in differentiation conditions for several days before ASCL1 or NEUROG2 expression (Masserdotti et al., 2015;Chouchane et al., 2017). This resistance to lineage conversion in astrocytes isolated at late developmental stages is likely explained by the epigenetic changes occurring in these cells during differentiation from an immature to mature state (Masserdotti et al., 2015). According to this interpretation, ASCL1-mediated lineage reprogramming of MGCs into iNs in adult mouse retina requires co-treatment with histone deacetylase-inhibitors (Jorstad et al., 2017).

Our data also showed that the presence of mitogenic factors EGF and FGF2 during MGC expansion substantially increased the efficiency of conversion into iNs, suggesting that exposure to those growth factors endows MGCs with higher plasticity. It has been shown that FGF2 induces methylation of Lysine 4 and suppresses methylation of Lysine 9 of histone H3 at the signal transducer and activator of transcription (STAT) binding site (Song and Ghosh, 2004), whereas EGF affects chromatin architecture at the regulatory element of cyclin D1, through a process involving Cre-binding protein (CBP), Histone deacetylase 1 (HDAC1) and Suv39h1 histone/chromatin remodeling complex (Lee et al., 2012). It is conceivable that such effects of EGF/FGF2 upon MGCs may facilitate the binding of NEUROG2 and ASCL1 to their target genes and, therefore, increases the efficiency of conversion. The effect was more pronounced for NEUROG2 than ASCL1; the latter has a known role in the remodeling of the chromatin landscape by itself, which increases its own accessibility to target genes (Raposo et al., 2015). In addition, expression of ASCL1 in NMDA-injured retinas of adult mice was sufficient to lineage reprogram only about 20% of MGCs into Otx2+ neurons, but this effect was enhanced by 2-fold in animals receiving concomitant treatment with trichostatin-A, a potent inhibitor of histone deacetylase (Jorstad et al., 2017).

Alternative explanations for the effects of EGF/FGF2 upon MGC-lineage conversion into iNs are rooted on the roles of these growth factors in cell proliferation (Todd et al., 2015) and survival (Rolf et al., 2003;Nickerson et al., 2011). In fact, we observed a larger fraction of NESTIN+, SOX2+ and BrdU+ cells in cultures grown in the presence of EGF/FGF2. However, such differences were much smaller than those observed in lineage-reprogramming efficiencies. Moreover, cell proliferation does not seem to be required for the conversion of astroglia into iNs (Heinrich et al., 2010;Chouchane et al., 2017). Survival of iNs appears to be a more important limiting factor during cell-lineage reprogramming (Gascon et al., 2016;Chouchane et al., 2017). However, we did not observe significant differences in cell death between MGCs expanded in the presence or absence of EGF/FGF2 using time-lapse video-microscopy (data not shown). Thus, the explanation that reprogramming is facilitated by EGF/FGF2 through effects upon chromatin structure remains a likely explanation, although further studies are needed to support this hypothesis.

A critical question in cell reprogramming is whether iNs acquire either single or multiple neuronal phenotypes (Amamoto and Arlotta, 2014;Heinrich et al., 2015). Recent work in our laboratory has shown that the origin of astroglial cells, either from cerebral cortex or cerebellum, affects the phenotype of iNs lineage-converted by either ASCL1 or NEUROG2 (Chouchane et al., 2017). Notably, most iNs generated from astroglia from either cerebellum or neocortex showed central hallmarks of neurons commonly observed in those regions, suggesting that a “molecular memory” in the astroglial cells contributes to the acquisition of specific neuronal phenotypes in iNs (Chouchane and Costa, 2018).

This notion is supported by our current finding that MGC-derived iNs upregulate several genes expressed in retinal neurons. Consistent with previous work (Pollak et al., 2013;Wohl and Reh, 2016;Jorstad et al., 2017), we found that ASCL1-converted MGCs generated iNs which express genes of horizontal and bipolar cells, but few genes commonly observed in photoreceptors, amacrine cells and RGCs, suggesting that these phenotypes are rare and/or incomplete in the converted cells. In contrast, expression of NEUROG2 converted MGCs into iNs accompanied by up-regulation of several genes of photoreceptors, amacrine cells and RGCs. Importantly, the latter phenotype is supported by the expression of two RGC-specific genes, namely Pou4f1 and Scl17a6 (Quina et al., 2005;Martersteck et al., 2017), as well as four other genes highly enriched in RGCs (Syn1, Parv, Calb2 and Tubb3). Immunolabeling confirmed the increased content of three of these markers (β_III_-TUBULIN, SYNAPSIN 1, PARVALBUMIN). Moreover, expression of the selective RGC marker RBPMS (Rodriguez et al., 2014) in MGC nucleofected with Neurog2 further suggests the acquisition of a RGC-like phenotype in iNs. These observations are particularly important, because previous attempts to lineage reprogram MGCs into iNs, both *in vitro* and *in vivo*, have failed to generate iNs with RGC-like phenotypes (Pollak et al., 2013;Wohl and Reh, 2016;Jorstad et al., 2017). Future experiments should address whether the expression of NEUROG2 in MGCs in the intact or injured adult retina are sufficient to lineage convert these cells into RGCs. Nevertheless, our observations in post-natal retinas indicate that NEUROG2 has an important instructive role for the RGC phenotype in vivo.

Altogether, our study corroborates previous evidence of the potential of MGC to reprogram into iNs following expression of ASCL1 (Pollak et al., 2013;Wohl and Reh, 2016;Jorstad et al., 2017), and provides compelling evidence that NEUROG2 expression is also sufficient to convert MGCs into iNs expressing several features of RGCs. Our results may thus contribute to develop new strategies of gene therapy aiming at the regeneration of retinal neurons in patients with glaucoma and other neurodegenerative retinopathies.

## Acknowledgements

We would like to thank Viviane Valença and Pedro Lucas França for their contribution with in vivo electroporation experiments. This work was supported by Conselho Nacional de Desenvolvimento Científico e Tecnológico (CNPq), Coordenação de Aperfeiçoamento de Pessoal de Nível Superior (CAPES), FAPERJ and INNT (Instituto Nacional de Neurociência Translacional).

## Legends to Supplementary Figures

**Fig S1. Polarized morphologies are more common in Neurog2- than Ascl1-iNs.**

Morphologies of iNs derived from MGCs nucleofected with either *Neurog2* or *Ascl1.* Cartesian plot representing the orientations of primary dendrites in iNs generated in different conditions (Neurog2 without GF = 31 iNs; Neurog2 with GF = 19 iNs; Ascl1 without GF = 21 iNs; Ascl1 with GF = 26 iNs). (C) Frequencies of iNs showing unipolar, bipolar or multipolar morphologies according to the TF used (Neurog2= 50 iNs; Asc1= 47 iNs). Observe the higher number of uni/bipolar among Neurog2-induced neurons (p<0.05; Chi-square test).

**Fig S2. Identification of major retina cell types using single-cell transcriptomes.**

(A) Clustering of 21,176 single-cell expression profiles into 39 retinal cell populations. The plot shows a two-dimensional representation (tSNE) of global gene expression relationships among 21,176 cells. (B) Annotation of the same cells into the 7 major retinal cell types, according to the expression of cell-type specific genes. (C) List of some differently expressed genes used to classify the 7 major retina cell types and also used to identify the phenotypes of iNs in this study (RT-qPCR and immunocytochemistry).

**Fig S3. Gene expression in cerebellar astroglia cells nucleofected with Neurog2 or Asc1.**

Graphic showing the relative expression levels (log_10_) of genes used to identify presumptive retina cell phenotypes (Figure 5) cerebellar astroglia cell cultures nucleofected with either Neurog2 (white bars) or Ascl1 (black bars). Observe that genes commonly observed in cerebellar neurons (Prox1, Vsx2, Slc32a1, Chat, Rbfox3 and Syn1) are upregulated, whereas genes whose expression is restricted to retinal neurons (Nrl, Rho, Pou4f1, Slc17a6) are not up regulated in cerebellar astroglia-derived iNs.

**Fig S4. Expression of** β**_III_-TUBULIN in RGCs generated in the postnatal retina following Neurog2-electroporation.**

(A) Coronal section of a P10 rat retina after electroporation with Neurog2-I-GFP at P0, immunolabeled for GFP (green) and β_III_-TUBULIN (TUBB3, red). Nuclei are stained with DAPI (blue). Image is a Z-projection of 8 confocal Z-stacks. Dashed box delimits a GFP+ cell within the ganglion cell layer (GCL). (B-C) Magnification of the dashed box in A showing the co-localization of GFP and β_III_-TUBULIN in a single confocal Z-stack. Scale bars: A: 50 µm; B-C: 25 µm.

**Legends to Supplementary Movies:**

**Supplementary Movie 1. MGC expanded in the presence of EGF/FGF2 and lineage reprogrammed into iNs by NEUROG2 show fast calcium transients**.

Movie shows 600 frames taken with 10ms exposure time and no interval. Observe the fast fluorescence intensity increase in the MGC-derived iN indicated by a red arrow in figure 4 A and B. MGCs in the same field show slow oscillations in fluorescence.

**Supplementary Movie 2. MGC expanded in the absence of EGF/FGF2 and lineagereprogrammed into iNs by NEUROG2 show fast calcium transients.**

Movie shows 600 frames taken with 10ms exposure time and no interval. Observe the fast fluorescence intensity increase in the MGC-derived iN indicated by a red arrow in figure 4 C and D. MGCs in the same field show slow oscillations in fluorescence.

